# Nullcline Analysis Provides Dynamic Mechanisms for the Differences in Electrical Activity of Distinct Subpopulations of Midbrain Dopamine Neurons

**DOI:** 10.64898/2026.02.02.703228

**Authors:** Christopher J Knowlton, Strahinja Stojanovic, Marle Jahnke, Jochen Roeper, Carmen C Canavier

## Abstract

Previously, electrophysiological differences between subpopulations of midbrain dopamine (DA) neurons were identified based on projection targets, including distinct responses to hyperpolarization and in the regularity of pacemaking. Here we explored single-compartment models of three subpopulations of DA neurons, projecting to medial shell of the nucleus accumbens (VTA-mNAcc), dorsomedial striatum (SNc-DMS) or dorsolateral striatum (SNc-DLS). We reduced the dimensionality to a phase plane consisting of membrane potential and one slow variable, either total slow potassium conductance or Kv4 channel inactivation. Nullclines are curves on which the rate of change of each variable is zero, given the value of the other variable. The voltage nullclines had three branches: upper spiking, unstable middle, and lower quiescent branch. Recruitment of Kv4 channels by the more prominent after-hyperpolarizing potential (AHP) in the DA-DMS and DA-DLS models channels stabilized pacemaking by creating a restorative moving fixed point along the quiescent branch. The slow inactivation of KV4 channels dominated and regularized the dynamics during the interspike interval; a dominant slow process may be a general mechanisn for stable regular pacemaking in a frequency range between 1-10 Hz. In contrast, the smaller AHP in VTA-mNAcc models prevented recruitment of this Kv4-mediated moving fixed point, which increased the sensitivity to synaptic inputs. On rebound from hyperpolarization the ability to produce robust ramps reverses between the DA neurons: now VTA-mNAcc projecting DA models fully recruited Kv4 channels and produced stable ramp-like pauses, whereas SNc-DLS projecting cells recruited significant regenerative inward CaV3 channels that overwhelmed Kv4 channels and produced ‘rebound’ bursts.

**Author Summary:** Midbraim dopamine (DA) neurons in the mammalian midbrain are linked to motivation, control of voluntary movement initiation, and reward-based learning. Their dysfunction is implicated in major disorders like Parkinson’s disease, schizophrenia or substance use disorders. Firing patters like bursts or pauses in most DA subpopulations are thought to signal better or worse than expected outcomes. Here we use dynamic systems analysis to capture how functional diversity of DA neurons of their intrinsic properties results in differences of synaptic input integration leading to the generation of burst and pause patterns of electrical activity.

## Introduction

Midbrain dopamine (DA) neurons can be differentiated in at least three major respects: the synaptic inputs they receive, their intrinsic dynamics, and in their axonal target projections. The focus of this study is how intrinsic dynamics differ between defined DA subpopulations, and how those differences are predicted to shape the integration of synaptic inputs and in turn the generation of behaviorally-relevant firing patterns such as bursts and pauses (1). Bursts in midbrain dopamine neurons have functional significance in boosting phasic DA release and stimulating D1-type DA receptors. Bursts are generally thought to depend on excitatory neurotransmission acting upon NMDA receptors (2). Pauses also have functional significance by reducing occupancy on D2 receptors in projection areas(1).

Here we differentiate three electrophysiological phenotypes based on the intrinsic responses of these subpopulations to excitation and hyperpolarization (Fig. 1B). Early studies focused on a generally more lateral, conventional population of DA neurons with projection targets in the dorsal striatum and the lateral shell of the nucleus accumbens (Fig 1A). These spontaneously pacing neurons (Fig. 1C2 and C3) exhibit very regular pacemaking, a pronounced after hyperpolarization (AHP), tall and wide (∼3 ms) action potentials with large spiking currents. They possess a limited dynamic range (<10 Hz) and a prominent voltage sag upon hyperpolarization (Fig. 1B). During these early studies, DA neurons were thought to be a homogeneous population incapable of firing faster than 10 Hz in response to injection of depolarizing current (3,4) without simultaneous NMDA receptor activation. Thus, bursts in DA neurons were originally operationally defined as beginning with an ISI <80 ms (5), corresponding to about 12 Hz, which exceeds the slow (1–10 Hz) single-spike firing rates usually observed *in vivo* (6).

**Figure 1:**
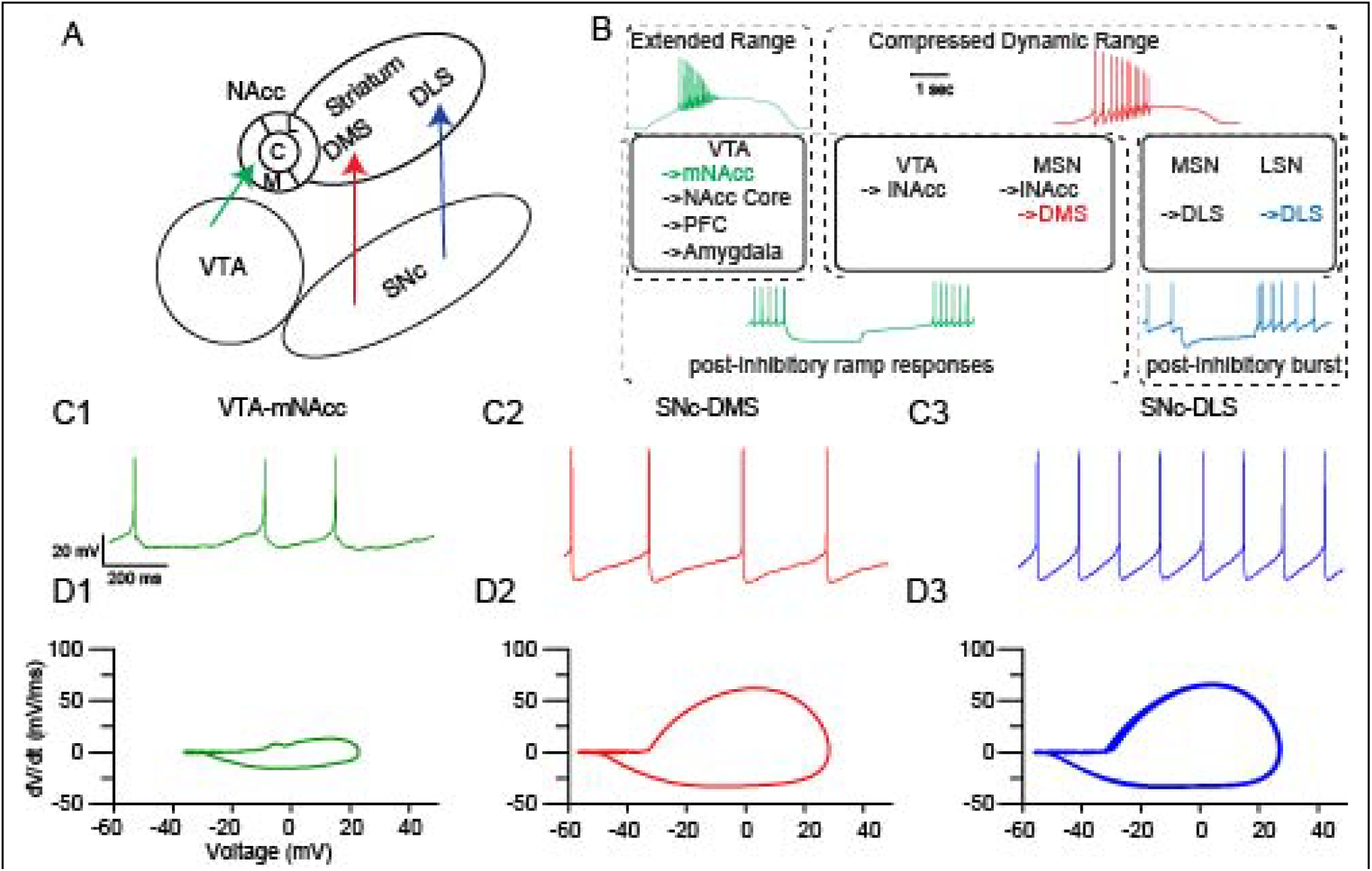
Subpopulation Electrophysiological Phenotypes with Exemplars. A: Schematic of a coronal slice of mouse midbrain (medial left, lateral right) with projection targets indicated by the arrow (->). Modelled subpopulation exemplars are the VTA-mNAcc (green), SNc – DMS (red), and SNc – DLS (blue). B. Schematic of relationship between projection target and electrophysiological phenotype. mNAcc projecting DA cells exhibit extended dynamic range (∼20 hz sustained) in response to a triangular waveform of depolarizing current, prolonged ramps after hyperpolarization, and a lack of sag potential. DMS projecting DA cells exhibit a compressed dynamic dynamic range (< 10 Hz) but also exhibit ramp responses after hyperpolarization (albeit short ramps than the mNAcc-projecting DA cells). DLS projecting DA cells also exhibit a compressed dynamic range but exhibit a unique rebound burst upon release from hyperpolarization. Model traces shown are color coded by projection target. Populations listed in black have the same phenotype as the color coded exemplar but are not illustrated. C. Representative Experimental Traces *in vitro*. D. Phase plane plots of experimental traces above.

In this study, we first differentiate two subpopulations based on responses to triangular ramps of excitation (top traces in Fig. 1B) the limited dynamic range does not apply to the more recently discovered atypical electrophysiological phenotype of dopamine neurons projecting ( Fig. 1A and B) to the prefrontal cortex, the medial shell and core of the nucleus accumbens, and other targets (7–11). Computational modeling (12) suggests that a lower availability of the FGF12 auxiliary subunit of the NaV channel that promotes long term inactivation (13,14) enables the higher frequency in this atypical, extended dynamic range population, which can fire in the “burst” range 12-20 Hz in response to somatic depolarization in the absence of NMDA receptor activation (9). Note that the atypical population has less regular pacemaking in vitro than the conventional DA subtypes (Fig. 1C1 versus C2 and C3). In this study, we provide a mechanistic explanation for the differences in regularity.

The conventional, compressed dynamic range DA phenotype can be subdivided further based on responses to hyperpolarization to form the three distinct electrophysiological phenotypes indicated by the solid rectangles in Fig. 1B. Most DA neurons respond to a hyperpolarizing step with a ramp in the membrane potential prior to resumption of pacemaking (Fig. 1B), with a longer ramp in the atypical compared to the conventional populations (15). However, a subset of the compressed dynamic range subpopulation, comprised specifically of DA neurons projecting to the DLS (Fig 1B bottom right) emit a burst of spikes with short latency after release from hyperpolarization (16). In this study we provide an explanation for the differential responses to hyperpolarization based on our dymamical systems analysis.

Defining the differential mechanisms that control DA subpopulation-specific intrinsic dynamics will not only provide a deeper understanding of their contribution to the behavioral dimension of dopamine in the context of motivated behavior but also facilitate the identification of causal drivers in the pathophysiology of DA-related psychiatric (e.g. schizophrenia, depression, substance use disorders, ADHD) and neurological disorders (e.g. Parkinson Disease, Tourette Syndrome).

## Computational Methods

### Computational Model Development

A source-target code (17) combines somatic location with axonal projection targets as shown in Fig. 1. We differentiate between two major phenotypes based on their response to excitation (9): a compressed dynamic firing range DA population (DA_CR_) and an extended dynamic range DA population (DA_ER_) (upper left vs upper right boxes in Fig. 1). We segregate two additional phenotypes based on post-inhibitory subthreshold properties (16): ramp response (DA_ramp_) and burst rebound (DA_burst_) phenotypes (lower left vs lower right boxes in Fig. 1). This anatomical source-target code can contribute complementary insights to those obtained from single-cell resolved transcriptomic studies of DA subpopulations (18–23).

Models of a representative neuron in each of the three DA subpopulations were created using the same currents shown in the equivalent circuit diagram in Fig. 2A but with differences in maximal conductances given in Table 1. One compartment models were chosen because, as we demonstrate below, they are amenable to mathematical analysis in the phase plane. The reduction of dimensionality required for this analysis is not feasible for a heterogeneous multicompartment model. We and others (12,24,25) have previously successfully used single compartment models to capture the dynamics of experimentally observed phenoma. In addition to the equivalent circuit, the cytosolic Ca^2+^ dynamics were taken into consideration to model the differential engagement of SK channels in the DA subpopulations. We assumed that the pool of Ca^2+^ sensed by the SK channel was a microdomain fed by nearby Ca^2+^ channels. Due to the limitations of a single compartment model, we assume that some fraction of Ca^2+^ entering via each Ca^2+^ channel contributes to this pool, whereas the remainder acts only electrogenically. The fractional Ca^2+^ coupling to each Ca^2+^ channel was based on experimental observations that the SK channel is primarily coupled to high threshold Ca^2+^ channels (represented by N-type Ca^2+^ channels in this model) in order to produce the AHP (26), with a contribution from T-type (16) but with limited coupling to the L-type (27) over pacing timescales. Ca^2+^ undergoes radial diffusion within the full diameter (1 μm) sized to represent a dendritic compartment and is dynamically buffered (28,29).

**Figure 2:**
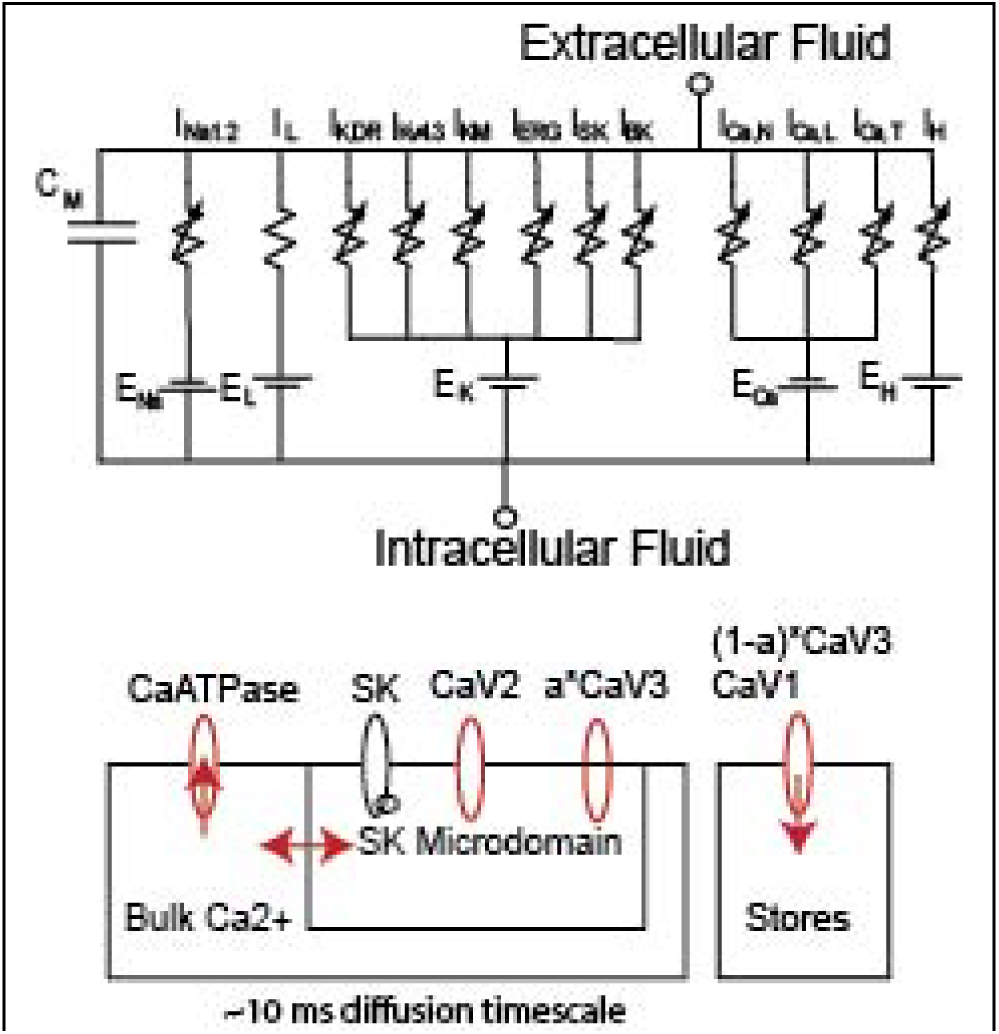
Model Schematic. A. Equivalent Circuit for Membrane Potential (see Methods). B. Calcium Handling (see Methods).

**Table 1:** Key Model Parameters. Representative models have varied parameters based on projection target. The Kv4.3 with a slow auxiliary subunit is present only in VTA-mNAcc model. CI1I2 is the transition rate into the long-term inactivation of NaV.

|  | VTA-mNAcc | SNc-DMS | SNc-DLS |
| --- | --- | --- | --- |
| gNaV (uS/cm2) | 7000 | 15000 | 15000 |
| gKdr (uS/cm2) | 1400 | 2000 | 2000 |
| tauKv4 (ms) | 44%: 65<br>56%: 150 | 75 | 35 |
| gCaT (uS/cm2) | 0 | 10 | 25 |
| gCaN (uS/cm2) | 200 | 200 | 200 |
| gCaL (uS/cm2) | 5 | 10 | 10 |
| gSK (uS/cm2) | 50 | 250 | 150 |
| gKv4 (uS/cm2) | 187.5 | 468.75 | 281.25 |
| gKM (uS/cm2) | 90 | 70 | 70 |
| gKERG (uS/cm2) | 5 | 10 | 10 |
| gHCN (uS/cm2) | 0 | 50 | 50 |
| gLNa (uS/cm2) | 3 | 6 | 4.5 |
| gLK (uS/cm2) | 5 | 9 | 9 |
| CI1I2 (/ms) | 0.05 | 0.2 | 0.2 |
| dCa (um) | 0.2 | 0.1 | 0.1 |

Population level differences in the calcium binding proteins (30) were neglected, thus the buffering capacity and rates of calcium binding were assumed to be constant across subpopulations. The shallower AHP amplitude in the atypical, limited dynamic range subpopulation is the single most reliable functional property for that distinguishes that subpopulation (9,31)). Accordingly, the model for the VTA-mNAcc projecting DA cells has a 3-5 fold lower maximum SK conductance and as well a larger micro-domain thus diluting the pool of Ca^2+^ seen by each SK channel. This corresponds to a putative larger mean separation between SK and Ca^2+^ channels. The atypical neu-rons located in the VTA also lack a subthreshold oscillation Ca^2+^ to recruit SK during pacemaking (32,33), thus the Ca2+ conductances for the L and T type conductances were also reduced in the medial shell projecting model..

The fast sodium channel responsible for action potential upstroke is implemented in a Markov model (12) that includes a long-term inactivation state. Action potential repolarization is achieved with a delayed rectifying potassium channel that qualitatively includes the effects of multiple fast activating potassium channels, such as BK and fast components of Kv4.3 that its HH formulation cannot adequately capture. The AHP is generated by a combination of SK, adapted from (34), and KM (Kv7) adapted from (35). The K_V_4.3 channel was adapted from (15,36), with KChip-subunits mediating fast (∼50 ms) and slow (∼200 ms) inactivation in the mNAcc projecting population. The DLS and DMS-projecting models assume that only KChip-subunits mediating fast inactivation are present. The relative values of AHP currents were chosen to replicate the effects of apamin (26) on AHP depth. The ERG channel (Kv11) (37) was included in all DA subtypes to regulate pacemaker frequency, particularly under simulated apamin, and allow for consistent repolarization from depolarized states under simulated spiking blockers. The K-ATP channel (38) was neglected since coupling between metabolic and electrical activity is not essential for the phenomena we model here. The N and L-type Ca^2+^ currents are included in all populations, with the N-type channel (CaV2) representing a composite of all high-threshold calcium channels (N, P/Q, R). The HCN (H current) (Neuhoff et al 2002) is modeled at electophysically calibrated levels in the conventional DLS and DMS-projecting model populations and omitted from the atypical medial shell projecting population. The model was developed in python and NEURON (39) and is freely available on Modeldb at (URL here) (reviewer password). Additionally, full model equations can be found in the supplemental materials.

### Fast/slow phase plane analysis using nullclines

The intrinsic dynamics of dopamine neurons can best be understood using a technique called separation of time scales, in which the state variables (Hodgkin-Huxley type gating variables and Markov model occupancy states) are subdivided into a fast and a slow group based on their dynamics (40). We collapse the dynamics of all the fast gating variables by setting them to their steady value as a function of membrane potential (and Ca^2+^ concentration where necessary) and track a single slow variable. This reduces the system to only two variables, membrane potential (V) and a much slower variable, which renders the system amenable to analysis in a phase plane (Fig. 3). The V nullcline (black curve) is the set of pairs of values of the membrane potential and the slow variable at which net current (which determines the rate of change) is zero, and the slow nullcline (green curve) is the set of pairs of values at which the rate of change of the slow variable is zero. The only possible resting potential is at the intersection of the two curves. However, the V nullcline for a pacemaker is nonmonotonic with three branches (41,42), and the intersection with the slow nullcline falls in the middle branch. The middle branch is created by the auto-catalytic, regenerative dynamics of currents like low threshold L-type CaV1.3 and the persistent NaV, modeled as residual occupancy in the open state in our Markov model. Positive feedback from regenerative currents destabilizes the resting potential as any slight depolarization opens channels and recruits more channel opening. Conversely, any slight hyperpolarization closes channels and causes more channels to close. Therefore, trajectories are pushed away from this branch (arrows). Under the fast/slow separation of variables, movement in the vertical direction is fast, toward the stable branches of V nullcline and away from middle branch; the dynamics evolve slowly along the two stable V nullcline branches. The lower stable branch represents a hyperpolarized branch in which the channels mediating the regenerative subthreshold currents are closed, whereas on the upper branch these channels are open. The direction of motion is determined by the slow variable, which decreases above and increases below the slow nullcline. Fig. 3 illustrates “relaxation oscillator” (43) dynamics in which the oscillatory trajectory (blue curve) follows the fast (V) nullcline and jumps between branches at the “knees”.

**Figure 3:**
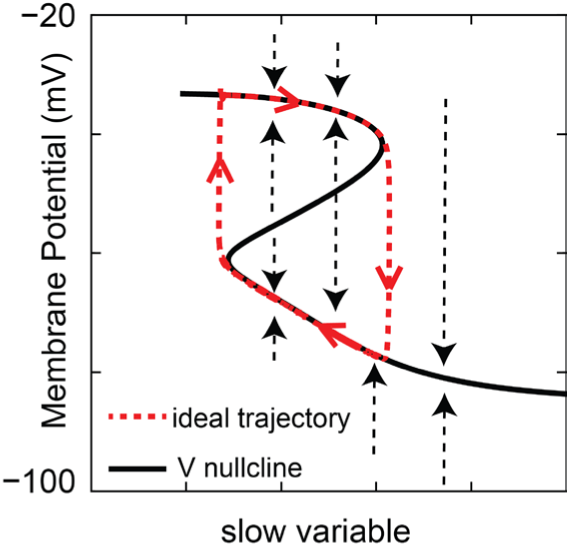
Fast/Slow analysis. The nullclines determine trajectories as shown by arrows denoting the vector field. Canavier et al., 2016

## Results

### Pacemaking

Two dimensional fast/slow dynamics provide useful insights but only apply approximately to DA neuron models because the upper branch of the V nullcline is not truly stable. Instead, it is a spiking branch because it is above the threshold for action potential generation. Technically, a third dimension orthogonal to the VmgKslow variable 2D-graph is required to allow the trajectory to spiral around the top branch during spiking as in (44). Figure 4 extends the simplistic model from Figure 3 to the more realistic model of a DMS projecting SNc DA neuron (SNc-DMS) using the channels from Figure 2A. Fig 4A shows a typical sequence of action potential (red), AHP (purple), and ramp (orange) under simulated *in vitro* conditions. The AHP is driven primarily by SK currents recruited by high threshold Ca^2+^ channels during the preceding spike (26). Figure 4B shows the voltage nullcline with respect to SK activation (black curve) along with the phase portrait of the dynamics (arrows in all panels indicate clockwise or counterclockwise directionality of the trajectory). The AHP dynamics are constrained to follow the stable lower branch; repolarization can only occur when the dynamics are released following the crossing of the knee at ∼-50 mV. Note that during the interspike interval (IS) indicated in orange, the dynamics are controlled by a different slow process, namely Ca^2+^ concentration, as illustrated in panel D and explained below. During regular pacemaking, the repolarization of a single action potential driven by the delayed rectifier (I_K,DR_) is sufficient to force a crossing of the unstable middle branch, resulting in a fast regenerative hyperpolarization towards the stable lower branch. As the separation between fast and slow dynamics? is relative, rather than absolute as it was in the toy model in Fig. 3, the dynamics will not follow the nullclines exactly. Note that due to the dynamics of SK activation by transient high threshold N-type Ca^2+^ currents, the steady state SK nullcline (and by extension the Ca^2+^ nullcline) is not informative and therefore not shown. Figure 4C shows the voltage vs total slow components of the potassium conductance (KV4, KV7, K,SK and KV11) in that phase space. The voltage nullcline was generated by finding the potassium conductance required to produce an outward current equal and opposite to the net inward steady state current of the model lacking those potassium conductances. This creates a source-agnostic nullcline for voltage with respect to potassium conductances. In contrast to Fig. 4B, in Fig. 4C, the trajectory closely follows the left knee of the nullcline particularly during the ramp-like interspike interval (orange), with the upstroke of the action potential occurring when the trajectory reaches the end of the stable lower branch, causing a jump at the knee between the stable lower and unstable middle branches. Note that despite the majority of time during pacing is occupied by a depolarizing ramp (orange), that phase occupies only a small segment around the lower knee of the voltage nullcline. To explore that region we assume that the slow potassium nullcline from Fig 4C is due entirely to Kv4.3 and that the activation gate of Kv4.3 is set to its steady state value for that voltage. Fig. 4D shows that the dominant slow variable during the ramp-like portion of the ISI is indeed the slow inactivation of Kv4, as the trajectory closely follows the channel between the steady state inactivation as a function of membrane potential (the nullcline for the inactivation variable dq/dt(V) = 0, dashed curve) and the voltage nullcline. Within this channel, perturbations in voltage are restored rapidly towards the voltage branch as Kv4 activation (and thus outward current) increases with depolarization and decreases with hyperpolarization, opposing the perturbation. Thus the dynamics are dominated by the slow inactivation of the Kv4 which causes a leftward movent on the yellow portion of the trajectory in Fig. 4D; inactivation decreases outward current causing an upward shift along the same part of the trajectory. The reason the movement is slow is because that rate of inactivation is given by the following equation (see full details in supporting information):

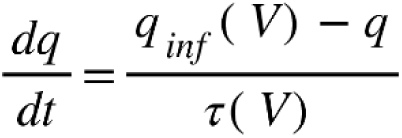

**Figure 4:**
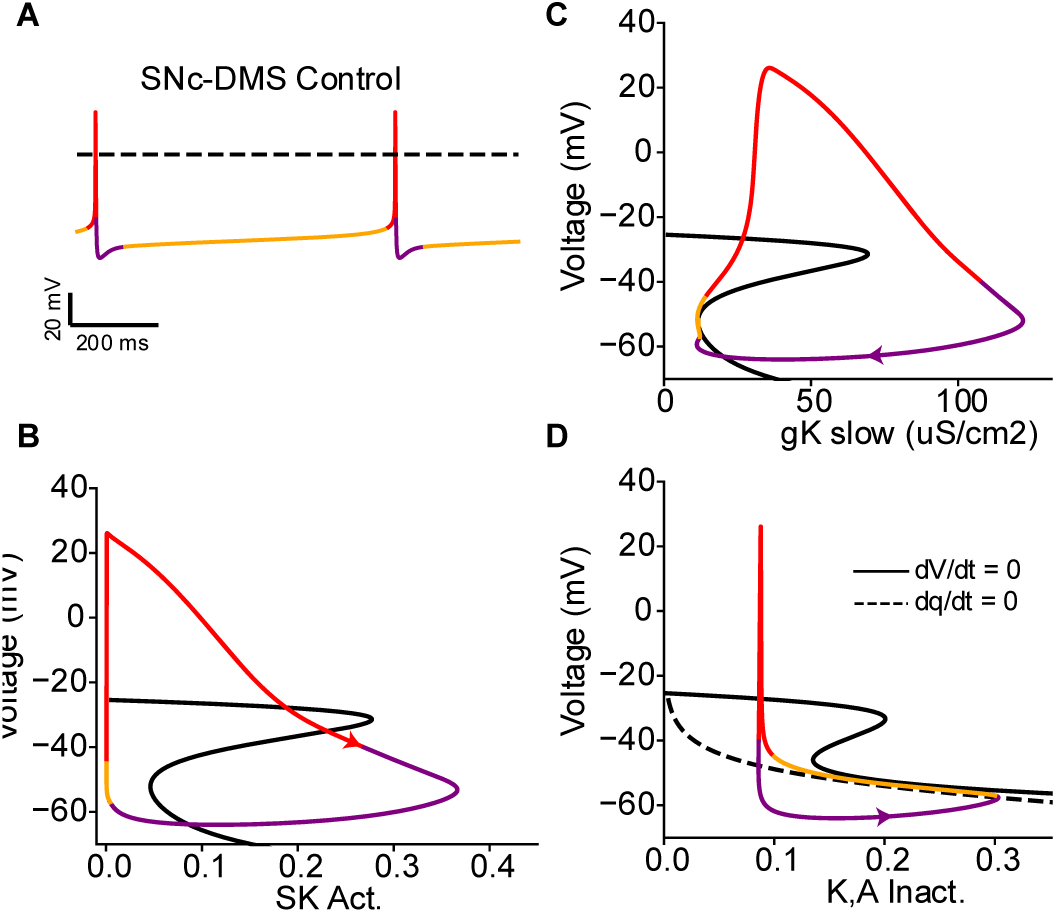
Fast/Slow analysis on SNc-DMS model. A: Voltage trace with color coded phases: action potential– red, AHP-purple, and ramp-orange. B: Phase space with respect to SK activation. C: Phase space with respect to total slow potassium (Kv7, Kv11, SK, Kv4). D: Phase space with respect to Kv4 inactivation.

In that equation, *q_inf_(V)* denotes the steady state value at a given membrane potential whereas *q* denotes the current value of the inactivation gate. The difference between these two values is small when the trajectory (q value) is near the steady state, and moreover the rate of change is obtained by dividing this small value by the relatively large (compared to the fast variables) inactivation time constant of ∼50 ms. Thus the Kv4 activation (via the voltage nullcline, soild line) keeps the voltage adjacent to the Kv4 inactivation nullcline (dashed line) while that adjacency limits the rate of change of the Kv4 inactivation variable itself – making the dynamics slower than the ∼50 ms time constant of inactivation would suggest. This tight proximity prevents the voltage dynamics from outpacing the slow Kv4 dynamics, only allowing regenerative depolarization after reaching the knee at the left end of the lower branch in Figure 4D.

An *in vitro* study of the regulation of pacemaking in VTA DA neurons focused selectively on the most regularly pacemaking neurons (45), which we now know were likely conventional neurons projecting to the lateral shell of the nucleus accumbens (9). That study showed that the interspike interval between two spikes in that population can be separated into the after-hyperpolarization (AHP) and a slow ramp leading to the next spike, as we have done in Fig. 4A. The AHP is initiated by the delayed rectifier but then sustained by the SK channel (46). The time course of the ramp in the DLS-projecting model is controlled by the slow inactivation of K_V_4 channels, as shown in Fig. 4D and predicted by (45). The depth of AHP forces the voltage trajectory of conventional neurons (multicolored trace in Fig. 4B-D) to approach, then move along, the lower branch of the V nullcline (black). The ramp results because the trajectory is confined to a narrow channel between the nullclines in which the rates of change of either variable is small (they are zero for each variable along its respective nullcline. On a timescale faster than the slow rates of change, we can envision a series of slowly drifting “resting potentials” (47) that are robust to noise perturbations in the vertical direction because convergence to the “pseudo” resting potential is fast compared to horizontal movement driven by slow inactivation of KV4.

Figure 5 contrasts conventional, limited dynamic range DMS-projecting model neurons with the atypical, extended dynamic range mNAcc-projecting model neuron. Figs 5A1 and B1 show representative model traces during pacemaking, with colored dots sampling the membrane potential every 100 ms during the interspike interval. Figs. 5A2 and B2 plot the pacemaking trajectory in the phase plane consisting of membrane potential and KV4 inactivation. In the DMS projecting model in Fig. 5A2, Kv4 availability increases (inactivation is removed) during the AHP as shown by the rightward motion below the nullclines to about-60 mV, reversing direction only after crossing the Kv4 nullcline as the AHP currents deactivate. The ramp response is confined to the region between the voltage and Kv4 nullclines, with points indicating 100 ms intervals during the ramplike ISI. The region between those nullclines can be treated as moving fixed point as shown in the instantaneous IV curves in Figure 5A3. In this case a moving fixed point corresponds to a resting potential; it is only truly fixed at a fixed value of the slow inactivation; since the value of the inactivation changes slowly, the “resting” potential moves slowly. We generated an instantaneous IV curve (42) by fixing the values of the slowest variables, consisting of Kv4 inactivation (both slow and fast subunits for VTA-mNAcc), HCN activation, CaV3 inactivation, SK activation, KV7 activation, and KV11 open fraction, at their values at the indicated points on the trajectory, then finding the steady state current at membrane potentials from-56 to +40mV given that set of fixed slow variables. The colored red dots on the trajectory (gray) are fixed points on a fast time scale because brief perturbations of the membrane potential in either direction are pushed back to those points by a restorative current proportional to the steep slope of the instantaneous IV curves. The slope is equal to the instantaneous restorative conductance. The evolution of the slow variables, primarily the inactivation of KV4 drives the movement of the rolling? Instantaneous fixed point. The instantaneous fixed point loses its influence on the dynamics when the Kv4 channel is replaced as the dominant channel by recruitment of regeneratively inward CaL and NaV channels at depolarized voltages.

**Figure 5.**
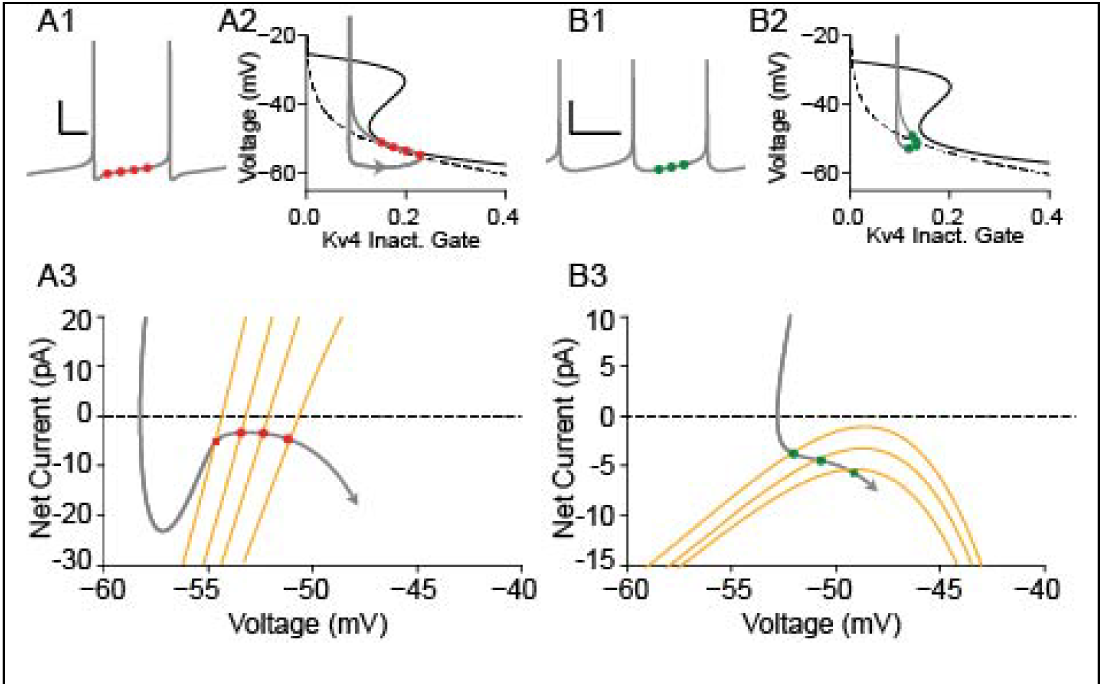
DMS, but not mNAcc projecting models are regular pacemakers. A: DMS model A1: Voltage trace of typical ISI. Dots indicate points in 50 ms intervals post-AP at which slow variables are sampled. A2: Voltage vs Kv4 inactivation phase space. Dashed line indicates dq/dt=0 while solid line indicates dV/dt=0 with all other variables at their steady state. A3: Subthreshold instantaneous IV curves (orange). Capacitive current (trajectory) vs voltage of trace in A1 and 2 is shown in gray. **B:** mNAcc projecting model. B1-B3: As in A1-3.

The single most reliable marker to discriminate between the conventional and atypical subpopulations is that the latter has a consistently shallower AHP (9,31), mediated by a smaller SK conductance putatively aided by weak coupling to fewer Ca^2+^ channels. Accordingly, modeling shows that the AHP and the ramp phases of the conventional population ISI (45), are driven by two different slow processes. During the AHP, calcium dynamics driving the SK current dominate and recruit Kv4, but the inactivation of KV4 dominates during the ramp phase. This recruitment does not happen in the atypical population due to the shallower AHP evident in the voltage trace in Fig. 5B1. In the mNAcc projecting model (Fig 5B), the moving fixed-point analogy does not apply. Despite having a similar narrow channel between the Kv4 inactivation and voltage nullclines, the lack of a deep AHP prevents substantial recruitment of Kv4 (Fig. 5B2). The lack of recruitment of Kv4 results in relatively flat instantaneous I-V curves (Fig. 5B3) during the ISI with small slopes, resulting in only weakly or nonexistent restorative Kv4-mediated currents.

The lack of strong restorative Kv4-mediated currents would predict that mNAcc projecting cells would be more sensitive to noise. Therefore we analyzed data from spontaneously active dopamine neurons identified by projection target, with representative examples shown in Fig 6A1 and A2. The atypical, medial shell projecting population did indeed have a larger coefficient of variation (CV) than the two conventional DMS and DLS-projecting population (Fig. 6B). Figure 6B shows that under control in-vitro conditions (see Methods) DA cells identified to project to the mNAcc from the VTA (6A1) have a larger coefficient of variation than those projecting to DMS or DLS (Fig. 6A2) with mNAcc: 40.46+/-18.18% N=20, DMS: 11.22+/-7.07% N=21, DLS: 12.13+/-8.05% N=17. To illustrate the mechanism, we applied a normal-distributed, mean zero noise sginal analogous to the intrinsic channel noise of recorded cells *in vitro*. In the DMS (and SNc-DLS, which is analogous to SNc-DMS in this regard but example traces not shown) models (Figure 6C3,4), perturbations during the ramp portion of the ISI remain confined within the region between the pair of attractive nullclines, rapidly restoring the dynamics to the mobile fixed point. In contrast, perturbations in the mNAcc projecting model (Figure 6C1,2) occur entirely in the unconstrained left section of phase space. In this region perturbations are not rapidly restored and can rapidly summate to produce rapid spiking. Additionally, the presence of the Kv4 nullcline allows for some hyperpolarizing transients to perturb into the region with slower dynamics – enhancing the delay the subsequent spike. Under increasing levels of additive noise, (Fig 6D), the coefficient of variation in the SNc-DMS and SNc-DLS models are significantly lower than that of the VTA-mNAcc models over levels consistent with realistic in-vitro environments.

**Figure 6:**
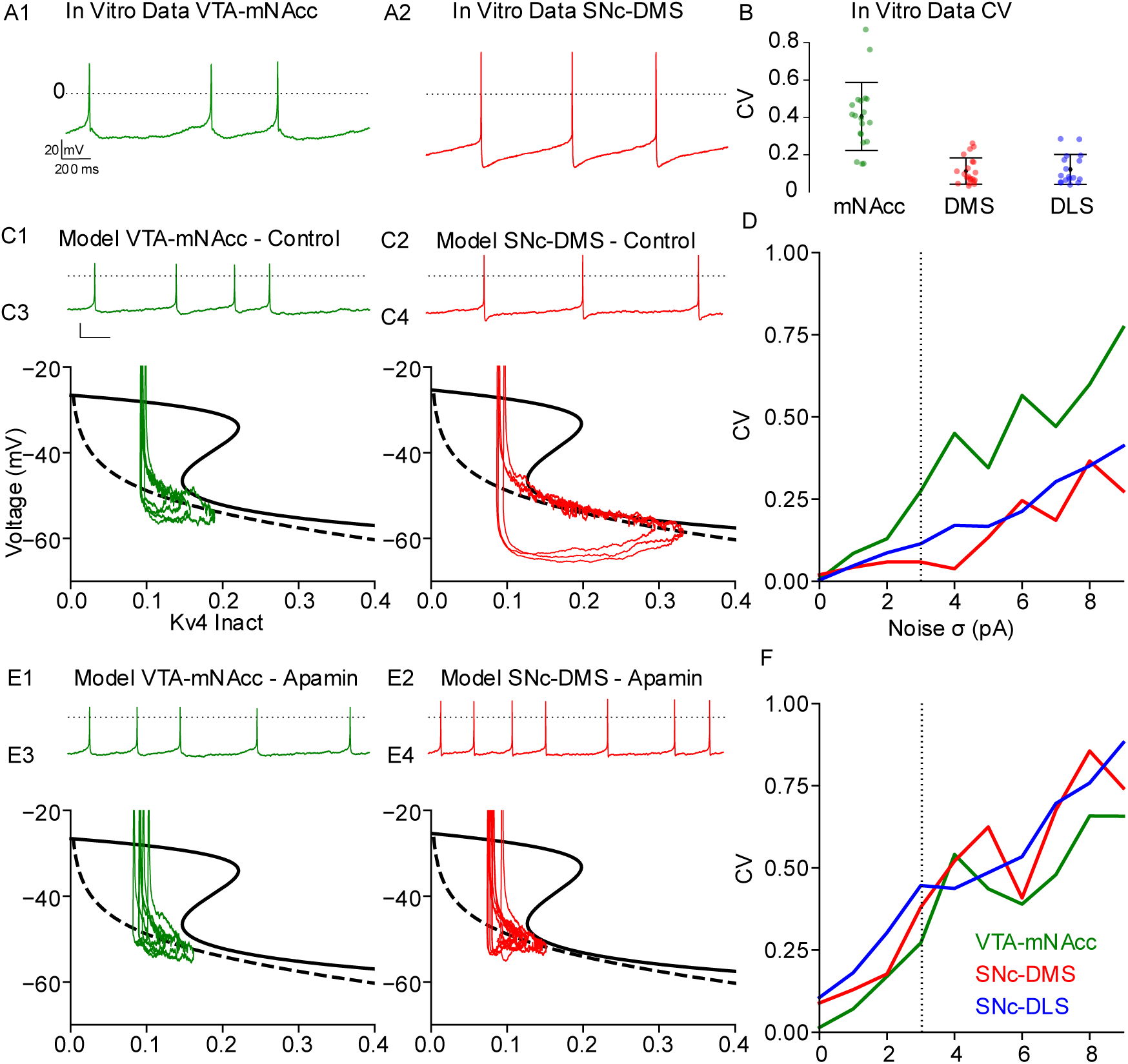
Mechanism for Robust Pacemaking in conventional but not atypical DA neurons. In Vitro figure A: Representative in-vitro traces for DMS, mNAcc projecting cells identified by retrograde tracing (see Methods). **B:** CV for projection defined cells in vitro. VTA-mNAcc: (green) m=40.46 std=18.18 N=20, DMS: (red) m=11.22 std=7.07 N=21, DLS: (blue) m=12.13 std=8.05 N=17. **C**: Model response of (C1) SNc-DMS model voltage vs time and (C2) voltage vs Kv4 inactivation phase-space under 3 pA gaussian noise (C3) VTA-mNAcc model voltage vs time and (C2) voltage vs Kv4 inactivation phase-space under 3 pA gaussian noise. **D:** CV of projection specific models vs mean synaptic noise input rate. DLS (blue) model examples are equivalent to DMS and not shown. **E:** Models from C under simulated apamin (gSK=0). **F:** CV for models under simulated apamin.

Setting the SK conductance to zero to simulate the bath application of apamin reduces the AHP and largely prevents the removal of Kv4 inactivation. Due to the lack of removal of inactivation, the dynamics of the DMS projecting model no longer occupies the stable region of phase space (Figure 6E1,2) and identical input pertubations evoke larger voltage excursions. The AHP contribution in the mNAcc projecting models is minimal and is predited to be largely unaffected by SK channels consistent with observed insensitivity to apamin among that population (48). As the control model does not remove significant activation of Kv4 during the AHP, a slight reduction in AHP would be predicted to have little effect on regularity. The coefficients of variation in the dorsal striatum projecting models are significantly increased by simulated SK block.

### Ramp versus Rebound Burst

The response to a step hyperpolarization to –80 mV (minimum of sag potential) for each of the projection specific models is shown in Figure 7. Consistent with recordings from identified subpopulations (9), and previous analyses (16) the mNAcc projecting DA model exhibits multi-second ramp responses (Fig. 7A1) and the DMS projecting DA model exhibits a ramp response of several hundred ms (Fig. 7A2), whereas the DLS projecting DA model exhibits a rebound burst (Fig. 7A3) with a significant reduction in the initial AHP. The differences in rebound responses can be attributed to the differences in the hyperpolarization activated currents. Figure 7B shows the intensive currents for CaV3 (purple), and Kv4 (teal) for the corresponding voltage traces in Fig 7A. The mNAcc projecting cell lacks significant contributions from the inward rebound channels HCN and CaV3, dominated exclusively by Kv4.3 (36). Additionally, the presence of the slow auxiliary subunit Kchip4a within this phenotype drastically increases the time constant of the Kv4 inactivation, leading to rebound delays significantly longer for that subpopulation over the nigral populations. During pacing (Fig 5B1-3), the regularity and ramp like response created by the dominance of the Kv4 channel was not recruited due to the lack of a significant AHP. Although pacemaking in the atypical VTA neurons does not exhibit a Kv4 driven ramp (49), a deep hyperpolarization evokes a Kv4 driven ramp because the hyperpolarization forces the trajectory into similar regions as traversed by conventional neurons during pacing. In response to hyperpolarizing steps or hyperpolarizing synaptic inputs (e.g. GABAB-R/D2R), Kv4 channels are significantly recruited, transiently allowing access to the ramp response.

**Figure 7:**
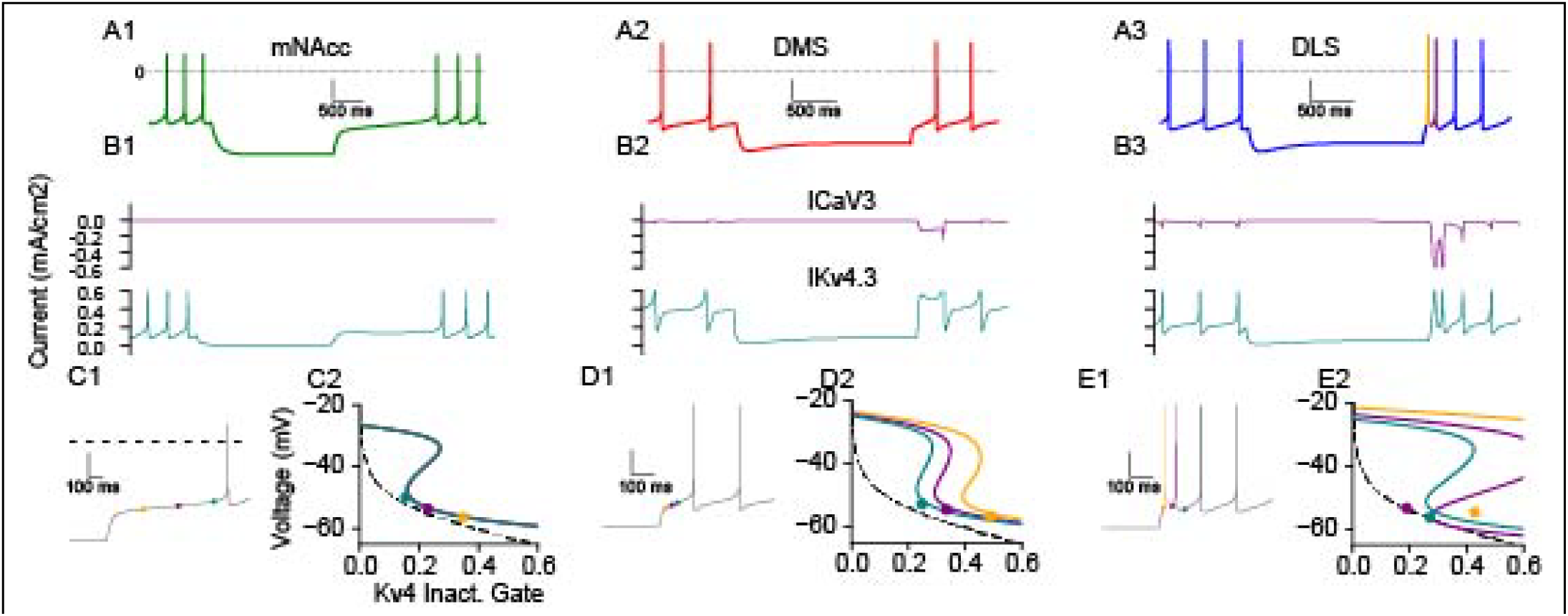
Rebound responses in mNAcc (left), DMS (center), and DLS (right) projecting models following 2s hyperpolarizations to-80 mV. A: Representative traces. Scalebars are 20 mV, 500 ms. B: Intensive CaV3 and Kv4 currents for traces from A. **C:** Rebound in mNAcc projecting model. C1: Zoom in on rebound from A1. Scale bars in C-E are 20 mV/100 ms. C2: Unlike during pacing (Fig 5B), dynamics are constrained by Kv4, voltage nullclines on rebound. **D:** Rebound in DMS projecting model. D1: Zoom in of A2. Colored points indicate points at which slow variables are sampled. D2: Instantaneous voltage nullclines at points in D1. Dynamics remains confined despite shifted nullclines. **E:** Rebound dynamics in DLS projecting model. E1: Zoom in of A2. First AP (orange) occurs between orange and purple points while 2nd spike (purple) occurs between purple and teal points. E2: Instantaneous voltage nullclines at points in E1. Larger CaV3 currents relative to Kv4.3 in DLS prevent eliminates confinement by Kv4.3 until inward rebound currents have weakened sufficiently.

Rebound bursts in the DLS projecting cells are consistent with CaV3 currents overwhelming Kv4 following deep hyperpolarization (16). While the mNAcc projecting cells only access the ramp phenotype during rebound from hyperpolarization, the inward calcium currents in DLS projecting cells overwhelms the Kv4.3 channels. Figure 7E shows both the steady state Kv4 nullcline along with the instantaneous voltage nullclines at three points, 40 (orange), 115 (purple), and 225 ms (teal) following the release of hyperpolarizing currents. The instantaneous nullclines were generated by fixing the other slow variables (SK activation, Ca_V_3 inactivation, HCN activation, ERG open fraction) at their current value. The inward currents shift the knee at which currents become regeneratively inward fully to the right of the dynamics until after the 2^rd^ AP (teal point). This temporary lack of confinement by Kv4 is due to strong transient inward currents from CaV3 and HCN. The combination of strong inward currents and the breakdown of the restorative mechanism allows for a period of significantly higher firing rates that might underlie the rebound bursts observed *in vivo* (50).

The DMS projecting DA cells have the full complement of rebound channels, but they notably have weaker CaV3 (51) and stronger Kv4 than DLS projecting DA cells. The presence of inward hyperpolarizing currents shifts the effective Kv4 voltage nullcline in a similar manner to the DLS projecting DA model (Figure 7D2); however, the regenerative region closes faster than the Kv4 inactivation gate closes. Thus at all points in the rebound ramp, the dynamics remain confined between the voltage and Kv4 nullclines.

## Discussion

### Robust slow pacemaking in the conventional population

Previous modeling work achieved the slow rates of rise during the ramp was by carefully balancing the dynamics of KV4 and the inward L-type CaV1.3 current (34). Here we showed that if the dynamics are constrained to a narrow region between the nullclines for voltage and inactivation of KV4, the balance between inward and outward currents does not need to be hand tuned to produce slow frequencies characteristic of convential DA neuron pacemaking. The nullcline for KV4 inactivation is simply the steady state inactivation curve as a function of membrane potential. The narrow region between the nullclines constrains the membrane potential to a ramp-like interspike interval that can be conceptualized as a slowly moving, pseudo “resting” potential. The evolution of the “resting” potential is driven by the time course of KV4 inactivation. An earlier study articulated the concept of a slowly moving fixed point (“resting” potential) but did not illustrate this concept (47). The key to the higher CVs exhibited by the atypical populations during pacemaking is their shallower AHP, which does not remove enough inactivation to effectively recruit KV4 that produces the ramplike ISI. The instantaneous IV curves in Fig. 5 reveal a much greater restorative slope conductance in the conventional (limited dynamic range) compared to atypical (etended dynamic range) models. Thus, we provide a mechanistic explanation for the higher CVs observed experimentally in the atypical populations compared to the conventional ones. Moreover, our results suggest that the 10% of the DA neurons that exhibited regular pacemaking activity *in vivo* (50), despite embedding within active circuits, were likely conventional DA neurons.

### Responses to Hyperpolarization

The VTA-mNAcc population exhibit more slowly inactivating Kv4 channels (52) and have a lower expression of hyperpolarization activated inward currents such as HCN or Ca_V_3. Although KV4 is not recruited during pacemaking in this population, hyperpolarizing steps in this population fully recruit Kv4.3 channels, providing robust pauses from relatively short hyperpolarizing steps (9,15). Modeling suggests that the ramp response to hyperpolarization in the atypical population results because the membrane potential is constrained in the same narrow channel between nullclines that causes the ramp in the interspike interval in the conventional population. The DLS and DMS-projecting populations exhibit higher levels of inward hyperpolarizing currents. During these transient driven hyperpolarizations, inward currents are recruited at a higher level in these population than KV4.3 – particularly in the DLS-projecting population with its larger CaV3 conductance (51). The instantaneous KV4 nullclines in Figure 7D suggest that ramp response to hyperpolarization in the DMS-projecting population is likely to be less robust than ramps during pacing. These results suggest that the rebound bursts observed *in vivo* (50) may be more likely to be emitted by DLS-projecting neurons (16). It is surprising and a bit counterintuitive that the difference in rebound response between DLS and DMS projecting DA neurons is not mediated by the H current, but this is consistent with experimental observations (16).

### Functional Implications of the Differences in How Subpopulations Integrate their inputs

Ramp-like responses in the mNAcc-projecting and to a lesser degree in the DMS-projecting DA neurons may amplify the response to inhibition. Stimulation of SNc dopamine neurons increases the probability of movement initiation (53,54). Thus the homogeneous responses of the DLS-projecting DA neurons to hyperpolarization may enable synchronized population rebound bursting in response to disinhibition that boosts DA release and thus envigorates movement initiation. In sum, the differences in excitability between the two populations are suggestive of intrinsic optimizations of their roles in the circuits in which they participate.

### Putative general principle for regular pacemaking

The dynamic mechanism for regularization of pacemaking by a slow variable described in this study may be a general biological organizing principle. For example, Fig. 1 of (55) shows that there is ramp phase in the pacemaking action potentials in the sinoatrial node of the rabbit. During this phase, as during the ramp phase of conventional, compressed firing range dopamine neurons, the net membrane current is very small suggesting the presence of sequential fixed points of the fast dynamics. As in this study, the sequence of “resting membrane potentials” may be driven by a slow variable.

A recent influential review (56) suggested that the mechanism for slow pacemaking in neurons is generally based on a steeply voltage-dependent persistent sodium current with a midpoint near−60 mV. Specifically for slow pacemaking in regularly firing midbrain dopamine neurons, the current required during the interspike interval is extremely small, ∼−1 to −5 pA (45), thus the net ionic current during that period is approximately the size of the current carried by a single ion channel (56). Given the stochasticity of channel opening, it is extremely unlikely that a single channel is responsible. Instead, it is instead likely that several macroscopic (both inward and outward) currents nearly cancel each other out during the ramp-like interspike interval, resulting in a net small inward current (57). Moreover, a previous computational study suggests that “achieving and maintaining a low firing rate is surprisingly difficult and fragile in a biological context(58)”. Thus, one commentary suggested that a small current carried by numerous membrane pores with a fS rather than a pS single channel conductance magnitude might be better suited to generate regular pacemaking (57). However, the mechanism that we suggest here with dynamics driven by a slow variable moving the trajectory through a channel near the voltage nullcline would remove the problem of precisely balancing the macroscopic inward and outward currents, favoring the hypothesis of balanced currents.

## Experimental Methods

All experimental procedures involving mice were approved by the German Regional Council of Darmstadt (V54-19c20/15-FU/1257).

### Methods – In Vitro Patch, Retrograde Tracing

C57BL/6N mice were anesthetized using isoflurane (AbbVie, USA; induction, 3.5%; maintenance, 0.8 to 1.4% in O2, 0.35 L/min) and placed in a stereotaxic frame (Kopf). A topical anaesthetic (lidocaine gel; EMLA crème, AstraZeneca, UK) was applied to the incision site. Throughout surgery, body temperature, respiratory rate (1–2 Hz), and reflexes were continuously monitored. Craniotomies were performed using a stereotaxic drill (0.5 mm diameter) to mNAcc (bregma: 1.54 mm, lateral: ±0.45 mm, ventral:-4.1 mm), lNAcc (bregma: 0.86 mm, lateral: ±1.75 mm, ventral:-4.5 mm), DLS + (bregma:, +0.74 mm; ML, 2.2 mm; DV, 2.6 mm), and DMS (bregma: +0.74 mm; ML, 1.2 mm; DV, 2.6 mm). Red retrobeads (100nl; Lumaflor) diluted (1:30) in artificial cerebrospinal fluid (ACSF; Harvard Apparatus) were injected either in the mNAc or lNAc using a 1 μL Hamilton syringe (Hamilton, Switzerland). Patch-clamp experiments were performed 2 to 4 days after tracer injection.

### Slice preparation

Mice were anesthetized by intraperitoneal injection of ketamine (250 mg/kg; Ketaset, Zoetis) and medetomidine hydrochloride (2.5 mg/kg; Domitor, OrionPharma) before intracardial perfusion using ice-cold ACSF (containing 125 mM NaCl, 2.5 mM KCl, 6 mM MgCl2, 0.1 mM CaCl2, 25 mM NaHCO3, 1.25 mM NaH2PO4, 50 mM sucrose, 2.5 mM glucose, 3 mM kynurenic acid, oxygenated with 95% O2 and 5% CO2). The brain was harvested and the midbrain was cut into 250 μm thick coronal slices using a vibratome (Leica VT1200S, Leica Biosystems, Germany). Before the experiment, slices recovered for 1h at 37°C in oxygenated ACSF (125 mM NaCl, 3.5 mM KCl, 1.2 mM MgCl2, 1.2 mM CaCl2, 25 mM NaHCO3, 1.25 mM NaH2PO4, 22.5 mM sucrose and 2.5 mM glucose).

### In vitro electrophysiology

For patch-clamp experiments, the slices were transferred to a temperature-controlled recording chamber maintained at 37 °C (Temperature Controller VI, Luigs & Neumann, Germany) and consistently perfused with ACSF at a rate of 2–4 ml/min. To block synaptic transmission, CNQX (20 μM, Biotrend), DL-AP5 (10 μM, Tocris), (-)-Sulpiride (0.15µM, Tocris), CGP (50 nM, Tocris) and Gabazine (SR95531 4 μM, Biotrend) were added to the ACSF.

Neurons were visualized using a light microscope (Axioskope 2 FS plus, Zeiss, Germany) equipped with an infrared light-source (SOLIS-850C, ThorLabs, USA) and a digital camera (CS505MUP1, ThorLabs, USA). Retrogradely traced neurons were identified by excitation of red retrobeads with a filtered LED lamp (SOLIS-1C, ThorLabs, USA, filtered at 546/12 nm). Labeled DA Neurons were recorded using borosilicate glass pipettes (3–4 MΩ, GC150TF, Harvard Apparatus, USA) filled with an internal solution (135 mM K-gluconate, 5 mM KCl, 10 mM HEPES, 0.1 mM EGTA, 5 mM MgCl2, 0.075 mM CaCl2, 5 mM ATP, 1 mM GTP, 0.1% Neurobiotin, pH 7.35, 290–300 mOsmol). Recordings were obtained using an EPC-10 patch-clamp amplifier (HEKA Elektronik) at a sampling rate of 20 kHz and a low-pass Bessel filter (5 kHz).

Data acquisition and analysis were performed using PatchmasterNext (HEKA Elektronik, Germany) and MATLAB (MathWorks, USA), and statistical analyses were conducted with GraphPad Prism 10 (GraphPad Software). Only spontaneously active midbrain dopaminergic neurons showing stable pacemaker firing were included for analysis. Neurobiotin-filled neurons were verified post hoc by immunohistochemistry.

### Immunohistochemistry

Midbrain slices and harvested forebrains were stored overnight at 4 °C in a fixative solution consisting of 4% paraformaldehyde and 0.15% picric acid in phosphate-buffered saline (PBS; pH 7.4). On the following day, the tissue was transferred to a storage solution containing 10% sucrose and 0.05% NaN₃ in distilled water, and stored at 4 °C. Forebrains containing the striatum were cut into 80 µm-thick coronal slices using a vibrating microtome (VT1000S, Leica Biosystems, Germany). Slices were washed in PBS (0.2 M, pH 7.4) and then incubated in a blocking solution (0.2 M PBS containing 10% horse serum, 0.5% Triton X-100, and 0.2% BSA) for 1 h for striatal slices and 2 h for thicker patch slices. Subsequently, slices were incubated overnight at room temperature in a carrier solution (0.2 M PBS containing 1% horse serum, 0.5% Triton X-100, and 0.2% BSA) with anti-tyrosine hydroxylase antibody (1:1000; Millipore, Germany). The following day, slices were washed in PBS and incubated overnight at room temperature in carrier solution containing goat anti-rabbit Alexa Fluor 488 (1:750; Invitrogen, USA) and streptavidin–Alexa Fluor 568 (1:750; Invitrogen, USA). On the third day, slices were washed in PBS, incubated in DAPI (0.2 µL/mL) for 5 min, mounted on glass slides using Vectashield mounting medium (Vector Laboratories, USA), and stored at 4 °C.

### Limitations of Computational Model

Single compartment models cannot account for distal integration. Thus, the spiking waveform in these models lacks a sharp shift in dV/dt at spike initiation due to the AIS. Whereas this difference is largely stylistic with respect to pacemaker firing, features such as ectopic spikes and spike shape may be highly sensitive to the location and size of the AIS.

The Kv4 channel used here is a Hodgkin-Huxley type model based on a fit to voltage clamp steps (15). The model predicts a substantial window current that produces a level of sensitivity of the frequency of the channel to Kv4.3 that is inconsistent with experiments in which that channel has been blocked(45). Additionally, the HH activation and final available pool at firing is insufficient to produce currents that have been shown to be responsible for up to 1/3 of the potassium currents during repolarization. Both discrepencies with recording suggest that a Markov model may be required to accurately represent this channel, but this presents complications as the inactivation and activation gates on which the steady state nullclines are generated are emergent rather than intrinsic properties of the channel. A possible future workaround to analyze a model with a Markov model of Kv4 would be to suspend inactivation transitions in both directions to generate voltage nullclines and instantaneous IV curves with respect to the noninactivated pool as in Figures 4 and 5.

Although this simple model includes a steady state Ca^2+^ concentration with respect to voltage, the limited interaction of SK with the continuously active CaV1.3 Ca^2+^ channels (26,27) results in bulk Ca^2+^ dynamics that are largely decoupled from the Ca^2+^-sensitive SK current. In the models used here the SK channel couples primarily to CaV2 and CaV3 channels with calcium extrusion tuned towards balancing transient currents rather than relatively small window currents.

Thus, the steady state Ca^2+^ as sensed by SK at all voltages is nearly zero. Unlike previous models (34) with continuously active coupling between SK and CaV1.3 during the ISI, the SK channel in the current models is only appreciably active during the AHP due to CaV2 activation and to CaV3 activation upon rebound from hyperpolarization.

